# Suppression of drug resistance reveals a genetic mechanism of metabolic plasticity in malaria parasites

**DOI:** 10.1101/155523

**Authors:** Ann M. Guggisberg, Philip M. Frasse, Andrew J. Jezewski, Natasha M. Kafai, Aakash Y. Gandhi, Samuel J. Erlinger, Audrey R. Odom John

## Abstract

In the malaria parasite *Plasmodium falciparum*, synthesis of isoprenoids from glycolytic intermediates is essential for survival. The antimalarial fosmidomycin (FSM) inhibits isoprenoid synthesis. In *P. falciparum*, we identify a loss-of-function mutation in *HAD2* (PF3D7_1226300) as necessary for FSM resistance. Enzymatic characterization reveals that HAD2, a member of the haloacid dehalogenase-like hydrolase (HAD) superfamily, is a phosphatase. Harnessing a growth defect in resistant parasites, we select for suppression of HAD2-mediated FSM resistance and uncover hypomorphic suppressor mutations in the locus encoding the glycolytic enzyme phosphofructokinase (PFK9). Metabolic profiling demonstrates that FSM resistance is achieved via increased steady-state levels of MEP pathway and glycolytic intermediates and confirms reduced PFK9 function in the suppressed strains. We identify HAD2 as a novel regulator of malaria parasite metabolism and drug sensitivity and uncover PFK9 as a novel site of genetic metabolic plasticity in the parasite. Our study informs the biological functions of an evolutionarily conserved family of metabolic regulators and reveals a previously undescribed strategy by which malaria parasites adapt to cellular metabolic dysregulation.

**IMPORTANCE:** Unique and essential aspects of parasite metabolism are excellent targets for development of new antimalarials. An improved understanding of parasite metabolism and drug resistance mechanisms are urgently needed. The antibiotic fosmidomycin targets the synthesis of essential isoprenoid compounds from glucose and is a candidate for antimalarial development. Our study identifies a novel mechanism of drug resistance and further describes a family of metabolic regulators in the parasite. Using a novel forward genetic approach, we also uncover mutations that suppress drug resistance in the glycolytic enzyme PFK9. Thus, we identify an unexpected genetic mechanism of adaptation to metabolic insult that influences parasite fitness and tolerance to antimalarials.

## Introduction

Malaria remains a global health threat, infecting 216 million people per year and causing nearly half a million deaths, mainly of pregnant women and young children (1). Resistance to current therapies has limited control efforts for malaria (2, 3). New drugs and a deeper understanding of drug resistance mechanisms are urgently needed.

Malaria is caused by infection with unicellular eukaryotic parasites of the genus *Plasmodium*. The species *Plasmodium falciparum* is responsible for most life-threatening malarial disease. As an obligate intracellular parasite of human erythrocytes, *Plasmodium falciparum* has unique metabolic features that may be exploited to discover new drug targets and develop new therapies. In the red blood cell niche, *Plasmodium* parasites are highly dependent on glucose metabolism. Infection with *Plasmodium* spp. results in a nearly 100-fold increase in glucose import in red blood cells (4–6). Despite these energy requirements, the parasite demonstrates little aerobic respiration via the TCA cycle. Instead, it relies on anaerobic glycolysis to produce ATP (7–10).

Besides ATP production, glucose also has a number of anabolic fates in *P. falciparum*. One such fate is the synthesis of isoprenoids. Isoprenoids are a large class of hydrocarbons with extensive structural and functional diversity (11). In the malaria parasite, isoprenoids perform several essential functions, including protein prenylation, dolichylation, and synthesis of GPI anchors (12–14). Despite this diversity, all isoprenoids are synthesized from a common five-carbon building block, isopentyl pyrophosphate (IPP). Evolution has produced two distinct routes for IPP synthesis: the mevalonate pathway, found in archaea, fungi, animals, and the cytoplasm of plants; and the methylerythritol phosphate (MEP) pathway, found in most eubacteria, plant chloroplasts, and apicomplexan parasites such as *P. falciparum* (15). Because it is both essential for the parasite and absent from the human host, the MEP pathway is a compelling target for antimalarial development. The antibiotic and antimalarial fosmidomycin (FSM) is a competitive inhibitor of the first committed enzymatic step of the MEP pathway, catalyzed by 1-deoxy-D-xylulose-5- phosphate reductoisomerase (DXR, E.C. 1.1.1.267) (16–18). FSM has been validated as a specific inhibitor of the MEP pathway in *P. falciparum* (19) and is a valuable chemical tool to study MEP pathway biology and essential metabolism in the parasite. In this study, we find that FSM is also a useful tool for probing glycolytic metabolism upstream of the essential MEP pathway.

Parasites are likely to control the proportion of glucose used for energy production versus production of secondary metabolites, such as isoprenoids. We previously used a screen for FSM resistance to identify HAD1, a metabolic regulator whose loss results in increased levels of MEP pathway intermediates and resistance to MEP pathway inhibition. HAD1 is a cytoplasmic sugar phosphatase that dephosphorylates a number of sugar phosphate intermediates upstream of the MEP pathway (20, 21). HAD1 belongs to the haloacid dehalogenase-like hydrolase (HAD) enzyme superfamily and more specifically, the IIB and Cof-like hydrolase subfamilies (22). While HADs are found in all kingdoms of life, HAD1 is most related to bacterial members of this superfamily (20, 23), which have been implicated in metabolic regulation, stress response, and phosphate homeostasis (24–28). However, most members of this superfamily remain uncharacterized.

In this study, we describe the discovery of HAD2, a second HAD family member in *P. falciparum*. We find that HAD2 is a cytosolic phosphatase required for metabolic homeostasis. Loss of HAD2 dysregulates glycolysis and misroutes metabolites toward the MEP pathway, conferring drug resistance. In our study, we harness a fitness defect in *had2* parasite strains to employ an innovative screen for suppression of drug resistance in the parasite. Selection for suppression of drug resistance identifies mutations in *PFK9*, which encodes the canonical glycolytic regulatory enzyme, phosphofructokinase. Reduction in PFK9 activity rescues the metabolic dysregulation in our resistant mutants and restores FSM sensitivity. Our unique approach thus reveals PFK9 as a site of exceptional metabolic plasticity in the parasite and uncovers a novel genetic mechanism by which *P. falciparum* malaria parasites may adapt to metabolic stress and drug selective pressure.

This article was submitted to an online preprint archive (29).

## RESULTS

### A FSM^R^ strain possesses a nonsense allele of *HAD2*, homolog of the MEP pathway regulator HAD1

The MEP pathway is responsible for the synthesis of the essential isoprenoid precursors isopentenyl pyrophosphate (IPP) and dimethylallyl pyrophosphate (DMAPP). This pathway is specifically inhibited by the antibiotic FSM (19, 30, 31). We previously generated *P. falciparum* strains resistant to FSM. Mutations in *HAD1* (PF3D7_1033400) cause the resistance phenotype in a majority of these strains (20). However, the genetic and biochemical basis of FSM resistance in strain E2 remained unknown. As previously reported, we find that E2 is less sensitive to FSM than its wild-type 3D7 parental line (Fig. 1A) (20). Strain E2 has a FSM IC_50_ of 4.8 ± 1.2 μM, significantly greater than that of its parent strain (0.9 ± 0.06 μM) (p≤0.01, unpaired Student’s t-test).

**Figure 1.**
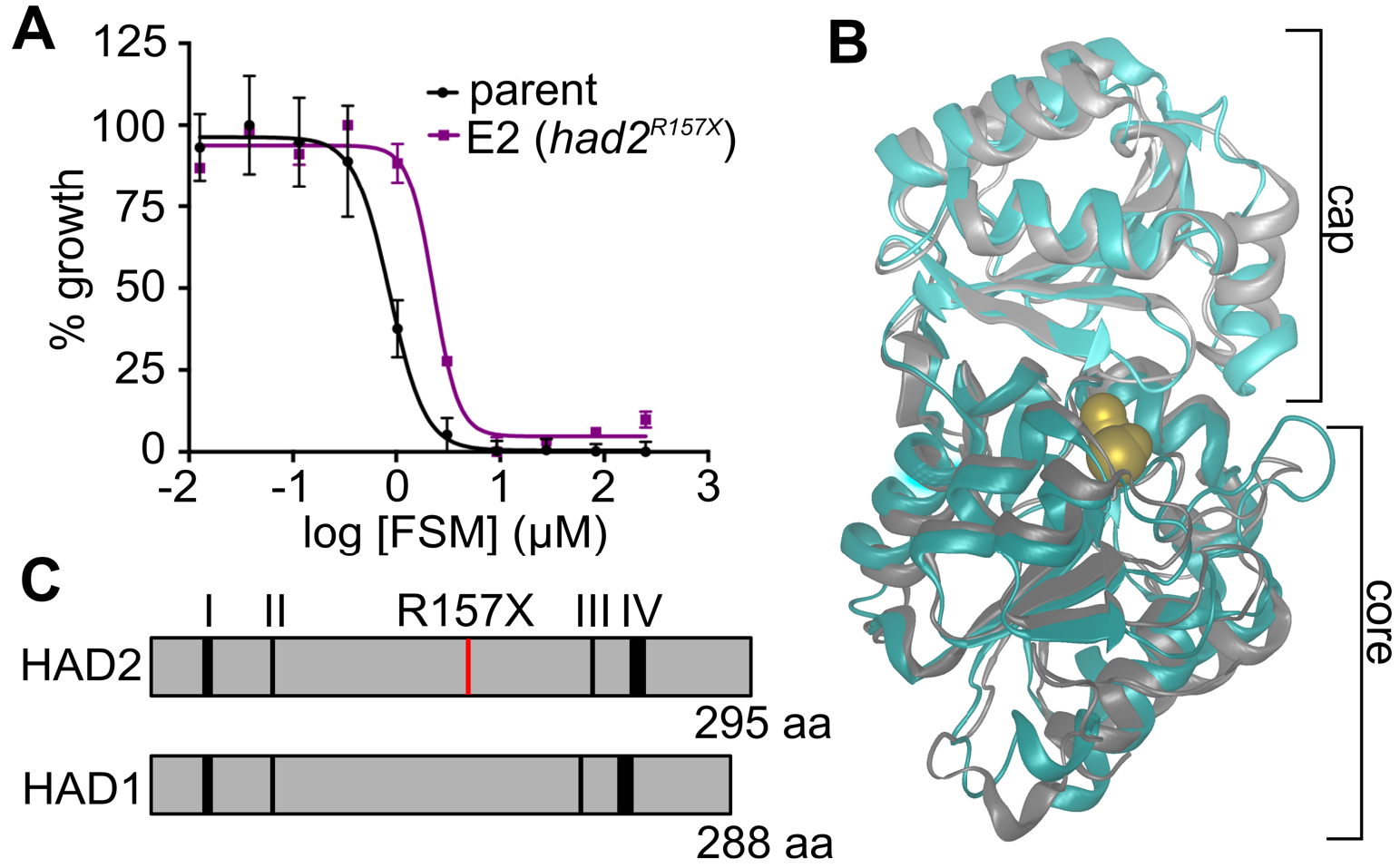
FSM^R^ strain E2 possesses a mutation in HAD2, a homolog of the MEP pathway regulator HAD1. **(A)** Representative FSM dose-response of the parental strain and strain E2. **(B)** *P. vivax* HAD2 (teal, PDB 2B30) is structurally similar to PfHAD1 (grey, PDB 4QJB). Ions (Mg^2+^, Ca^2+^, Cl^−^) are shown in yellow. **(C)** HAD2 is a homolog of HAD1 (29% identity and 53% similarity) and possesses all conserved HAD sequence motifs required for catalysis (37).

We find that this resistance phenotype is not due to changes in expression of the genes encoding the first two, rate-limiting, steps of the MEP pathway, *DXS* and *DXR* (32–35). In addition, strain E2 does not have genetic changes in the known FSM resistance locus and MEP pathway regulator, *HAD1*, nor are there changes in HAD1 expression by immunoblot (Figs. S1A and S1B).

To identify new genetic changes that may result in FSM resistance, we performed whole genome sequencing on strain E2 and identified an A469T mutation in PF3D7_1226300 (PlasmoDB ID), hereafter referred to as *HAD2* (36). Variant data for strains sequenced in this study can be found in Dataset S1. Sanger sequencing of the *HAD2* locus in strain E2 confirmed the presence of the A469T SNP. The A469T single nucleotide polymorphism (SNP) yields a truncated (R157X) protein variant and therefore we expect HAD2 function is lost in strain E2. Interestingly, HAD2 is a close homolog of the known MEP pathway regulator, the sugar phosphatase HAD1 (20). Sequence homology places both proteins in the haloacid dehalogenase-like hydrolase (HAD) superfamily and further within the IIB and Cof-like hydrolase subfamilies (Interpro IPR006379 and IPR000150, respectively) (22). While no structural information exists for *P. falciparum* HAD2, the structure of the *Plasmodium vivax* HAD2 (PVX_123945, PvHAD2) has been solved (PDB ID 2B30). PvHAD2 (93% identical and 98% similar to PfHAD2) contains the common structural motifs found in other HADs, including a core and cap domain (Fig. 1B). HAD2 possesses the four conserved sequence motifs found in HAD proteins (Fig. 1C), which are involved in binding of the substrate, coordination of the phosphoryl group and Mg^2+^ ion, and hydrolysis of the substrate phosphate (37–39). Overall, HAD2 and HAD1 protein sequences share ~29% sequence identity and ~53% sequence similarity (Fig. 1C). We hypothesized that HAD2, like HAD1, regulates metabolism in *P. falciparum*, and that loss of HAD2-mediated metabolic control was responsible for FSM-resistance in malaria parasite strain E2.

### HAD2 is a functional phosphometabolite phosphatase

We have previously established that *P. falciparum* HAD1 is a promiscuous sugar phosphatase, with activity against a wide range of phosphometabolites. Similarly, *P. vivax* HAD2 has been enzymatically characterized and found to possess phosphatase activity against various monophosphorylated substrates, including glycerol 2-phosphate (glc2P) and pyridoxal phosphate (PLP) (40). Recombinant PvHAD2 also utilizes additional monophosphorylated substrates, such as adenosine 5’-monophosphate (AMP) and glycerol 1-phosphate (glc1P), with moderate activity.

Based on the previous characterization of a close *Plasmodium* homolog, as well as sequence homology to HAD1 and other HAD proteins, we predicted that PfHAD2 would also function enzymatically as a phosphatase. We successfully purified recombinant PfHAD2 in *E. coli* and confirmed the phosphatase activity of recombinant PfHAD2 using para-nitrophenyl phosphate (*p*NPP), a promiscuous, chromogenic phospho-substrate (Fig. 2A) (23, 41). Because *E. coli* expresses a number of HAD-like phosphatases (23), we confirmed that the phosphatase activity was specific to purified PfHAD2 by expression and purification of a catalytically inactive mutant (HAD2^D26A^). The Asp26 residue was chosen for mutagenesis as the corresponding residue in PfHAD1 (Asp27) has been previously shown to be required for catalysis (21).

**Figure 2.**
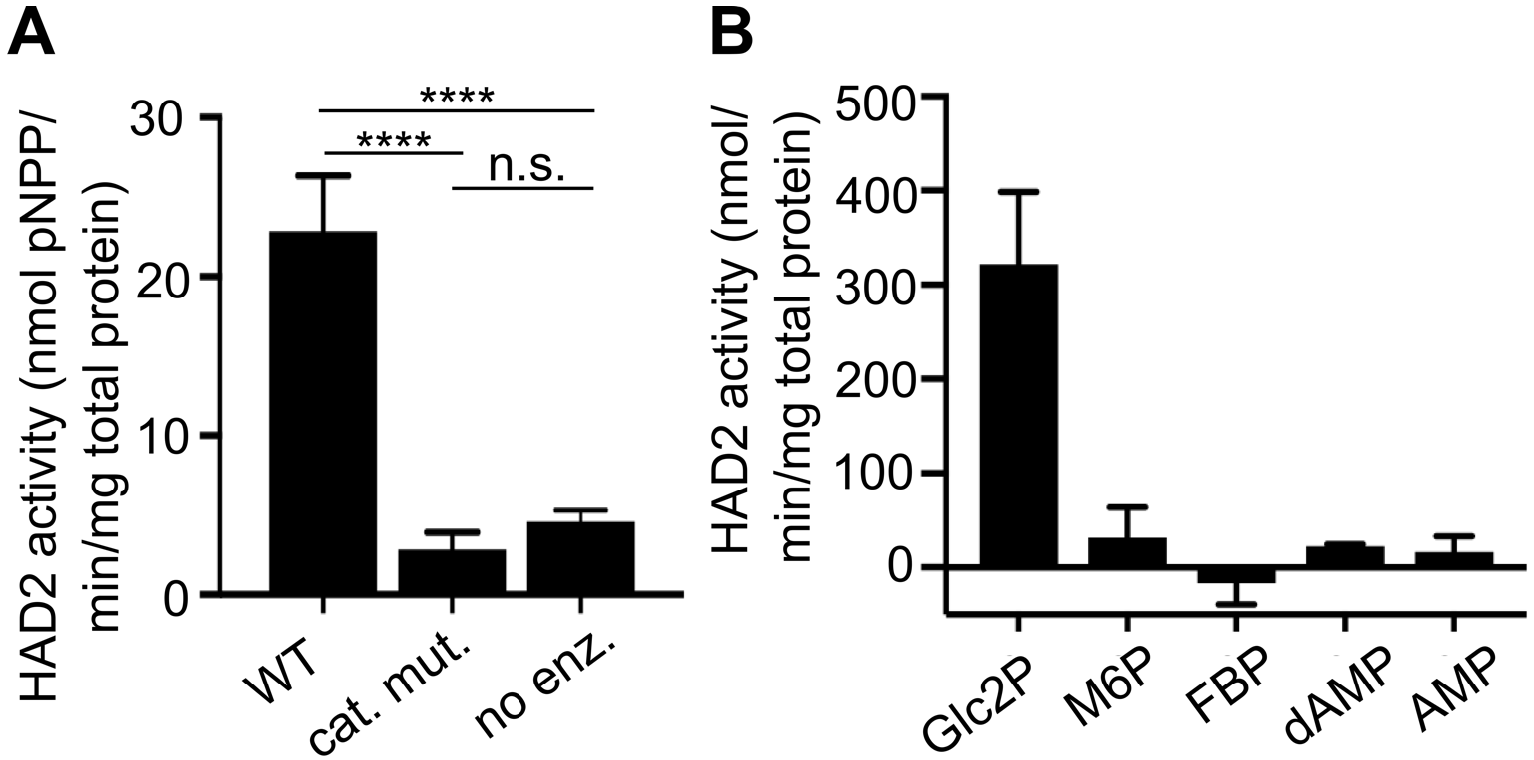
PfHAD2 is a phosphatase. **(A)** HAD2 is an active phosphatase, and HAD2^D26A^ is a catalytic mutant (cat. mut.) that can be used as a negative control for HAD2 specific activity. “No enz” represents a no enzyme control. Data shown are the enzyme activities using the synthetic phosphatase substrate *p*NPP. Error bars represent S.E.M. (**** = p≤0.0001, unpaired t-test). **(B)** Activity of HAD2, normalized to the activity of the catalytic mutant (HAD2^D26A^), for variety of substrates (2-GlcP, 2-glycerol-phosphate; M6P, mannose-6-phosphate; FBP, fructose-2,6-bisphosphate; dAMP, deoxy-adenosine monophosphate; AMP, adenosine monophosphate). Error bars represent S.E.M.

We also established the activity of PfHAD2 against a panel of phosphorylated substrates and determined that its substrate profile closely mirrors that of PvHAD2 (Fig. 2B). Overall, we find that PfHAD2 is a phosphatase with activity against small phosphosubstrates, such as *p*NPP and glc2P. These data suggest that, like HAD1 and related HADs in microbes and plants (23, 42–44), HAD2 is a promiscuous phosphatase with a preference for a variety of monophosphorylated phosphometabolites.

### *In vitro* evolution of mutations suppressing FSM resistance

During routine culturing of E2 FSM^R^ parasites, we observed that the E2 strain is growth-attenuated compared to its parental parasite strain. Surprisingly, during prolonged culture in the absence of FSM, this growth phenotype resolved, and improved growth rates correlated with a return to FSM sensitivity (Fig. 3A). From these observations, we hypothesized that *had2*^*R157X*^-mediated FSM resistance led to a fitness cost in cultured parasites. We sought to harness this fitness cost to drive *in vitro* evolution of a FSM-sensitive (FSM^S^) population, possessing additional, novel mutations that might suppress FSM resistance in *had2*^*R157X*^ parasite strains.

**Figure 3.**
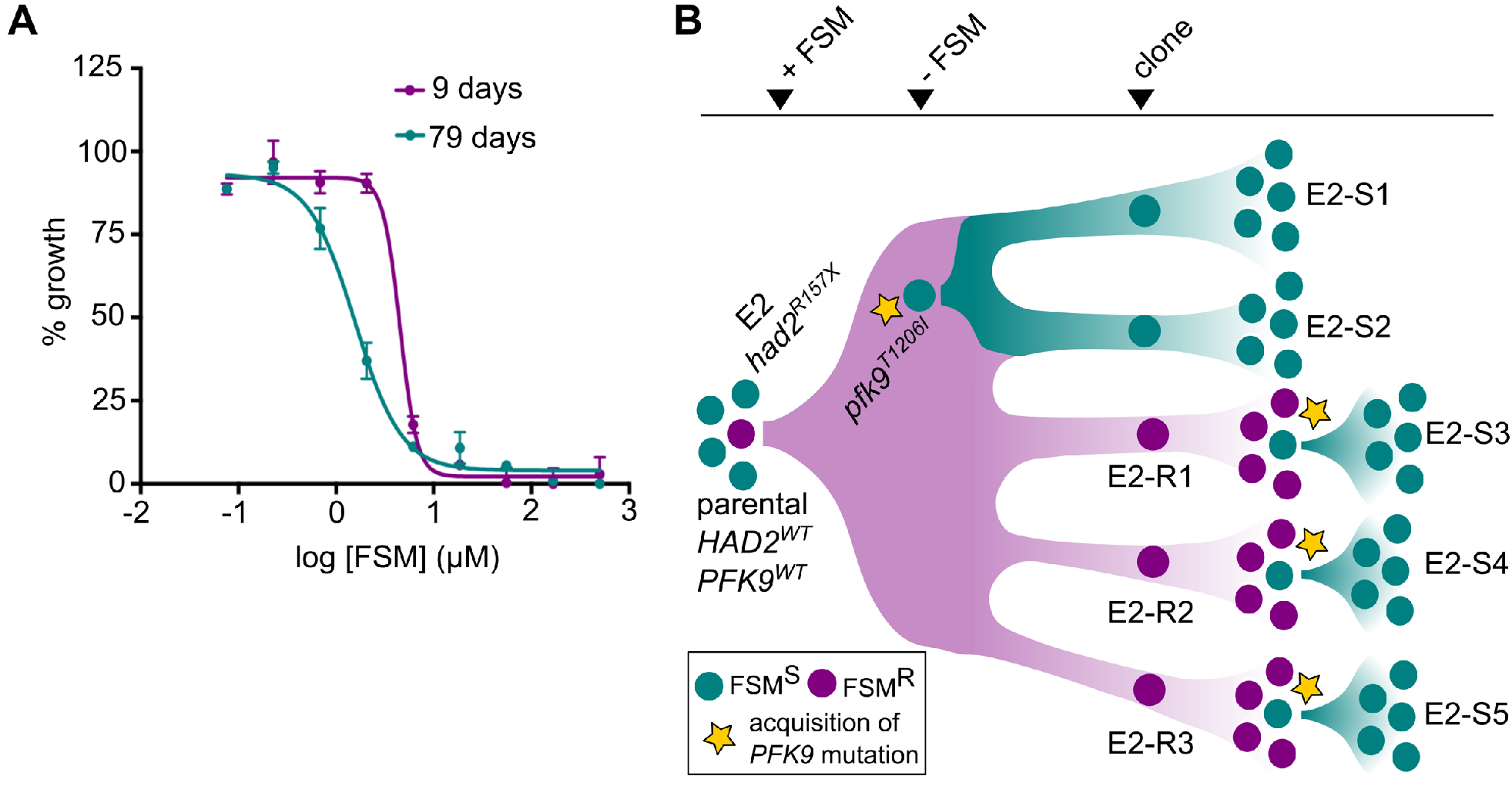
Leveraging a resistance-associated growth attenuation to identify genetic changes that modulate FSM sensitivity. **(A)** Prolonged culture results in loss of FSM resistance in strain E2. Shown are single FSM dose-responses of the strain E2 before (9 days) and after (79 days) prolonged culture without FSM. Nine days after thawing the resistant E2, we observe a FSM IC_50_ of 4.9 μM, while after 79 days of culture without FSM, E2 has an IC_50_ of 1.3 μM. Each data point is representative of the mean from two technical replicates. Error bars represent S.E.M. **(B)** Parasites are colored by their FSM phenotype (teal, FSM^S^; purple, FSM^R^). Cloned strains are named by FSM phenotype (E2-SX, sensitive; E2-RX, resistant). A FSM^S^ parental strain was selected under FSM pressure to enrich for FSM^R^ strain E2 (*had2*^*R157X*^). After relief of FSM pressure, a fitness advantage selects for spontaneous suppressor mutations in *PFK9* (*pfk9*^*mut*^, yellow star) that result in FSM sensitivity. FSM^R^ clones are grown without FSM pressure and a fitness advantage again selects for suppressor mutations in *PFK9* that result in increased growth rate and loss of FSM resistance.

FSM-resistant strain E2 was cultured through multiple passages in the absence of FSM selection. Through limiting dilution, we derived five E2-based clones in the absence of drug pressure (Fig. 3B). Three of the five clones remained FSM^R^ (designated clones E2-R1, -R2, and -R3), but two of these were found to be FSM^S^ (designated E2-S1 and -S2) (Fig. 3B and 4A). To validate our novel suppressor screen approach, we independently repeated this genetic selection with the three FSM^R^ E2 clones by again culturing in the absence of FSM for >1 month (Fig. 3B). As before, these strains (E2-S3, -S4, and -S5) also lost their FSM resistance phenotype (Fig. 4A).

**Figure 4.**
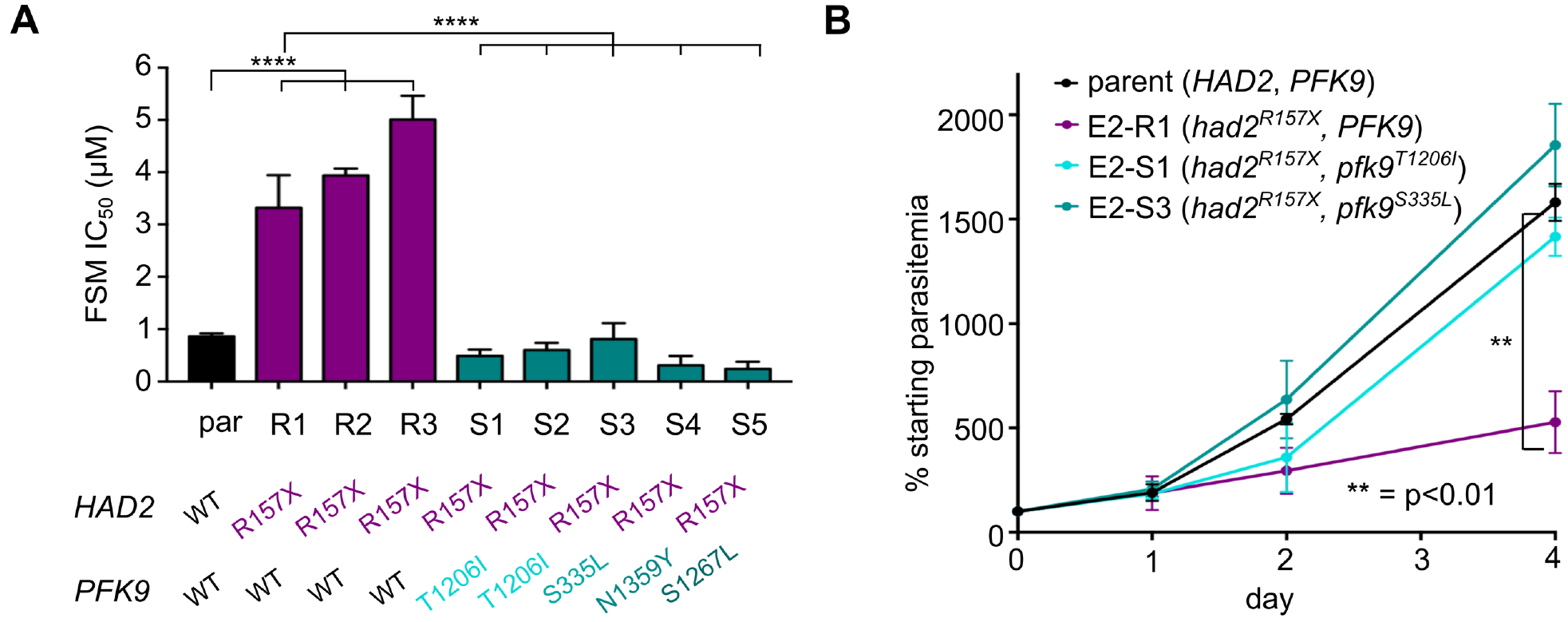
Suppressor strains with *PFK9* mutations display changes in FSM tolerance and growth. **(A)** Suppressed clones have significantly lower FSM IC50S (**** = p≤0.0001). Error bars represent S.E.M. *HAD2* and *PFK9* genotypes for each strain are indicated. **(B)** FSM resistance results in a fitness cost. A representative FSM^R^ clone with the *had2*^*R157X*^ allele (R1, purple line) has a reduced growth rate compared to the wild-type parental strain (black) (** = p≤0.01). The growth defect is rescued in two representative clones with mutations in *PFK9* (S1 and S3, teal lines). Growth is normalized to parasitemia on day 0. Error bars represent S.E.M. from independent growth experiments.

Consistent with our initial observation that our *had2*^*R157X*^ FSM resistant strain grows poorly, we find that the FSM^R^ clone E2-R1 grows at a significantly reduced rate compared to the parental strain, while two FSM^S^ clones (E2-S1 and E2-S3) have restored growth rates, similar to that of wild-type (Fig. 4B).

Loss of FSM resistance might have occurred by reversion of the *had2*^*R157X*^ mutation in E2-derived strains. Instead, we find that all E2-SX clones maintained loss of HAD2 via the *had2*^*R157X*^ mutation. We hypothesized that the FSM^S^ E2 clones, driven by a fitness advantage, had acquired new suppressor mutation(s) at an additional locus. We performed whole genome sequencing on the original five E2 clones to identify any genetic changes that segregated with FSM-sensitivity. Sequencing revealed that a new mutation (C3617T) in the locus encoding phosphofructokinase-9 (*PFK9*, PF3D7_0915400) is present in both suppressed (FSM^S^) E2 clones and none of the three FSM^R^ E2 clones (Fig. 3B, Dataset S1). The C3617T mutation results in a PFK9^T1206^ protein variant. *PFK9* contained the only SNP that segregated with the change in FSM tolerance. Two other loci had indels that also segregated with our FSM phenotype. These loci encode a tyrosine recombinase (MAL13P1.42) (45) and an erythrocyte surface protein (PIESP1, PFC0435w). Given their predicted functions and the presence of A/T indels in poly-A and poly-T tracts, we concluded that mutations in these loci were unlikely to result in our suppressed phenotype and prioritized *PFK9* as the likely locus of our suppressor mutation.

To verify whether mutations in *PFK9* were responsible for suppressing FSM resistance in all of our suppressed strains, we investigated *HAD2* and *PFK9* in the E2-S3, -S4, and -S5 strains, which were derived through independent evolution of the E2-R1, -R2, and -R3 populations in the absence of FSM. By Sanger sequencing, we find that, as before, all strains maintain the *had2*^*R157X*^ mutation and acquire new, independent *PFK9* mutations (Fig. 4A). The independent acquisition of four different *PFK9* alleles during selection, each of which was associated with both improved growth and loss of FSM sensitivity, strongly indicates that loss of PFK9 function is responsible for these phenotypes in strains lacking HAD2.

### Loss of HAD2 is necessary for FSM-resistance in *had2*^*R157X*^ parasites

HAD2 was not the sole genetic change in FSM^R^ strain E2. In addition, because intraerythrocytic *P. falciparum* parasites are haploid, we cannot distinguish recessive from dominant or gain-of-function mutations. Therefore, we sought to establish whether restoring HAD2 expression in *trans* in a *had2*^*R157X*^ strain would restore FSM sensitivity. Using a previously described expression system enabled by the piggyBac transposase (20, 46), we expressed HAD2-GFP driven by the *heat shock protein 110* (*Hsp110*) promoter (47). We confirmed that the transfected *had2^R157X^* E2-R2 clone maintains the *had2*^*R157X*^ allele at the endogenous locus and successfully expresses HAD2-GFP (Fig. 5A). Expression of HAD2-GFP in *had2*^*R157X*^ parasites results in restoration of FSM sensitivity (Fig. 5B), confirming that loss of HAD2 is necessary for FSM resistance in this strain. The resistant clone (E2-R2) has a FSM IC_50_ of 3.9 ± 0.2 μM, significantly higher than that of the wild-type parent strain (0.9 ± 0.06 μM, p≤0.001, one-way ANOVA and Sidak’s post-test). Expression of HAD2-GFP in E2-R2 results in an IC_50_ of 0.6 ± 0. 02 μM for FSM, significantly lower than the E2-R2 strain (p≤0.001) but not significantly different than the parental strain (p>0.5).

**Figure 5.**
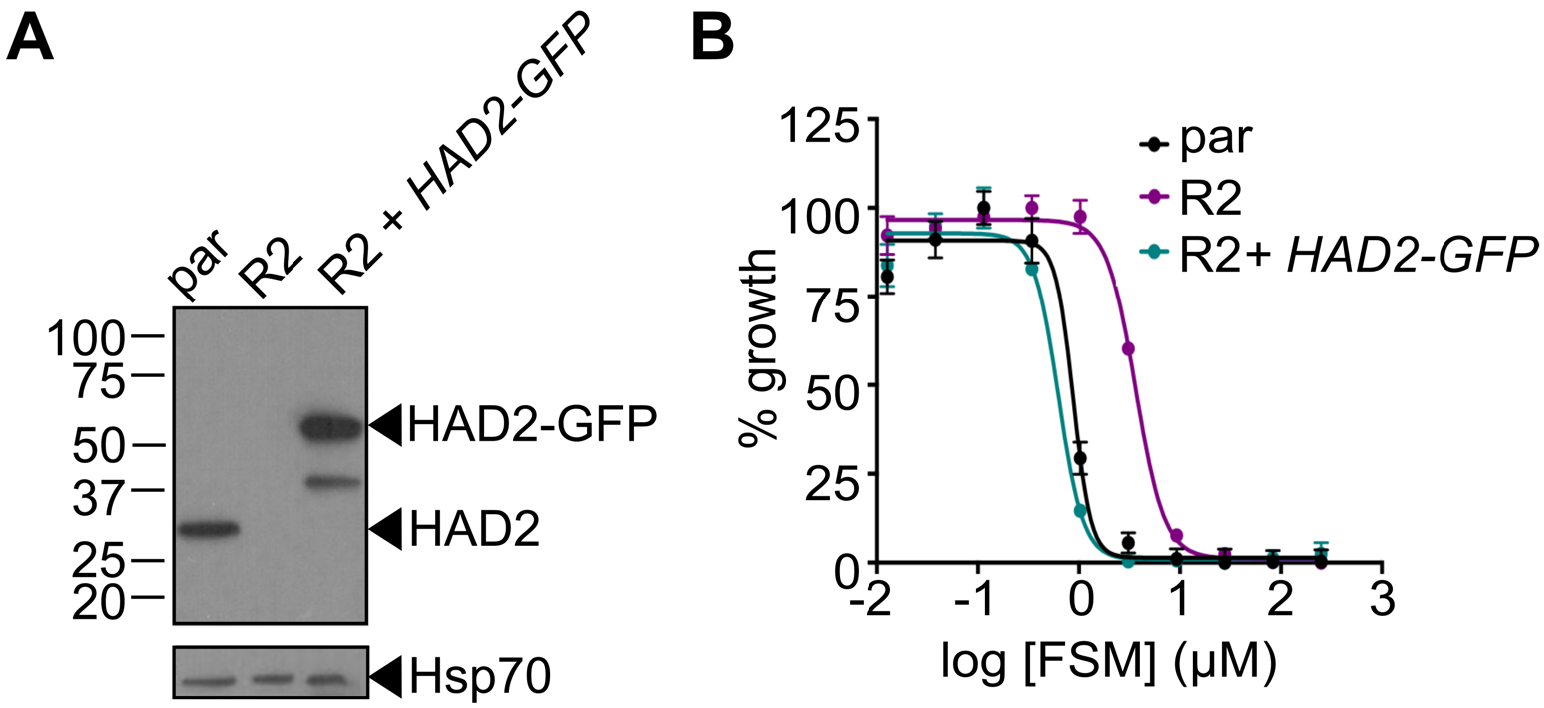
Loss of HAD2 is necessary for FSM resistance. **(A)** Successful expression of pTEOE110:HAD2-GFP in strain R2 (*had2*^*R157X*^, *PFK9*) was confirmed by immunoblot. Marker units are kilodaltons (kDa). The top blot was probed with anti-HAD2 antiserum (expected masses: HAD, 33 kDa; HAD2-GFP, 60 kDa). The bottom blot was probed with anti-heat shock protein 70 (Hsp70) antiserum as a loading control. **(B)** Representative FSM dose-response demonstrating expression of HAD2-GFP in strain R2 (*had2*^*R157X*^, *PFK9*) results in restored sensitivity to FSM. Strain R2 has an elevated FSM IC_50_ compared to the parental strain. When HAD2 expression is restored in strain R2, the resulting strain has an IC_50_ near that of the parent strain.

Using fluorescence microscopy, we also investigated the localization of HAD2-GFP in our E2-R2 Hsp110:HAD2-GFP strain. We observe that HAD2-GFP is diffusely present throughout the cytoplasm in asexual *P. falciparum* trophozoites and schizonts but excluded from the digestive food vacuole (Fig. S2). This finding is consistent with the lack of a predicted signal sequence for HAD2 using SignalP, PlasmoAP, and PlasMit algorithms (48–50).

### PFK9 mutations in suppressed strains are hypomorphic

The *PFK9* locus encodes the enzyme phosphofructokinase (PFK, E.C. 2.7.11), which catalyzes the first committed and canonically rate-limiting step of glycolysis, which is the conversion of fructose 6-phosphate to fructose 1,6-bisphosphate. PFK9 is comprised of two domains, alpha (α) and beta (β), which are typically encoded by independent genes in non-apicomplexans (51) (Fig. 6A). While in other systems the α domain is regulatory, previous work on *P. falciparum* PFK9 has demonstrated catalytic activity for both domains (51–55).

**Figure 6.**
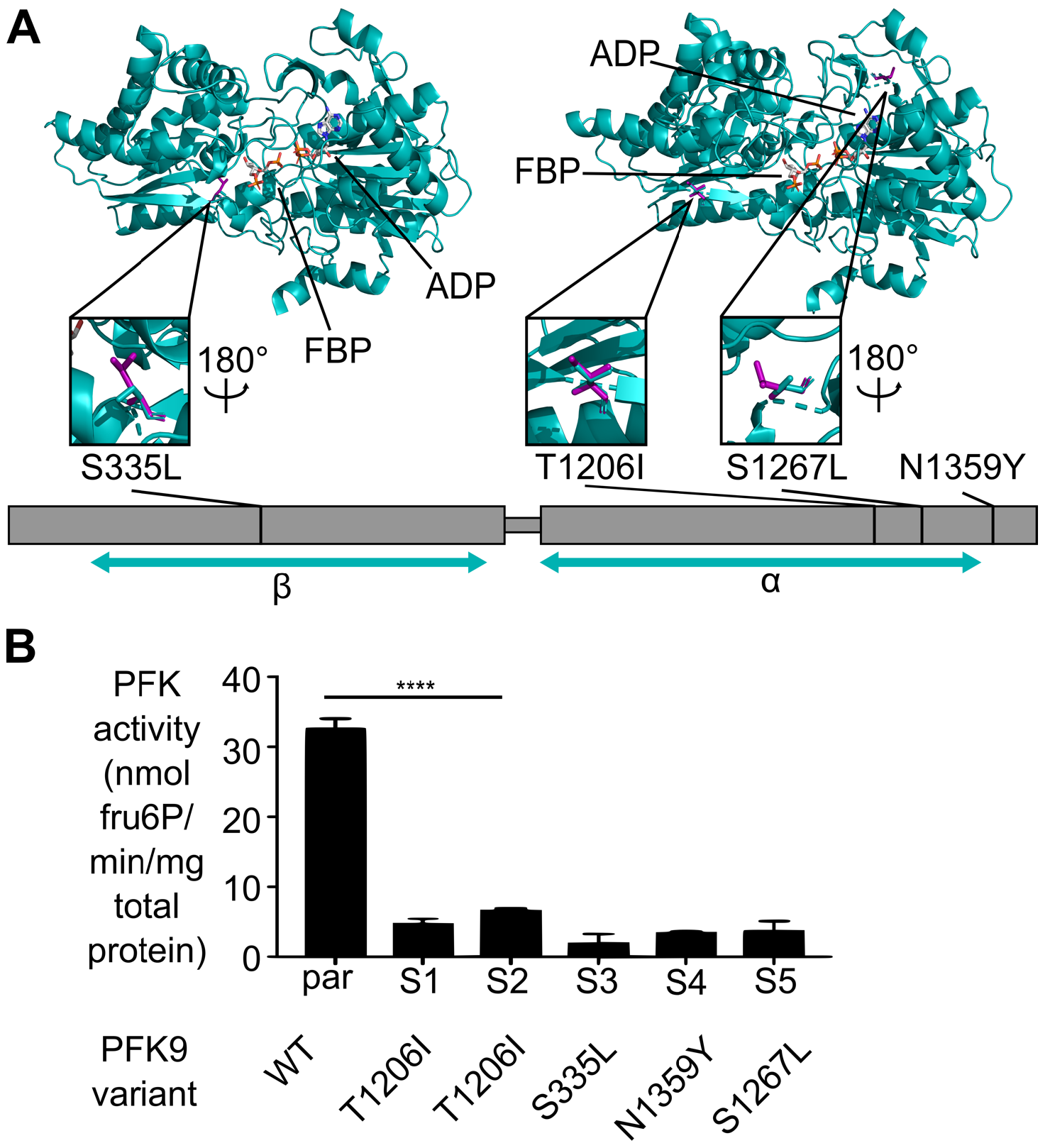
*PFK9* alleles in suppressed strains are hypomorphic. **(A)** Schematic of suppressor mutations found in *PFK9*. Resulting amino acid changes and strain names are indicated. Three of the four mutations are found on the structural model of PfPFK. The parts of the protein represented by the model are notated by the teal arrows under the α and β domains. Total protein length is 1418 amino acids. N1359Y fits outside of the model. The other mutations are represented by their stick model structure with the resulting change shown in magenta. Orientations of the close-up representations of the mutations are indicated when they differ from the main model. **(B)** Measurement of PFK activity of *P. falciparum* lysate indicates that E2-SX clones with *PFK9* suppressor mutations have significantly reduced PFK activity (**** = p≤0.0001, ANOVA, Sidak’s post-test). Error bars represent S.E.M. Assay is linear with respect to protein content and specific for PfPFK9 activity (S3 Fig).

Of the four PFK9 variants identified in this study, three variants map to the α domain, while one variant (S335L) maps to the β domain (Fig. 6A). We projected our mutations on a three-dimensional model of PfPFK9 to reveal a possible structural basis for altered PFK function. Three variants (S335L, T1206I, and S1267L) align and model to currently available crystal structures of PFK, while a fourth allele (N1359Y) does not. While three out of four mutations map to the α domain of PfPFK9 (T1206I, S1267L, N1359Y), these mutations do not appear to cluster in any particular region. All mutations affect amino acid residues that are physically distant from the substrate-binding pocket of either domain and are not predicted to impact binding or specific catalytic residues.

Consistent with a previous study on PfPFK9 (51), attempts to purify recombinant full-length PFK9 were unsuccessful. To then understand the enzymatic impacts of our PFK9 variants, we quantified the native PFK specific activity in *P. falciparum* (51, 56) (Fig. 6B, Fig. S3). Lysates from strains possessing *PFK9* mutations (E2-SX strains) have markedly reduced specific activity of PFK compared to the parental strain (Fig. 6B). Combined with the diverse mutation locations (Fig. 6A), the reduced lysate PFK activity in E2-SX strains suggests that a variety of genetic strategies to alter PFK function can lead to resistance suppression.

### Metabolic profiling reveals mechanisms of resistance and suppression in HAD2 and PFK9 mutant parasites

Reduced activity of PFK9, which catalyzes the canonical rate-limiting step in glycolysis, is associated with restored FSM sensitivity to *had2* mutant strains. Therefore, we anticipated that metabolic changes might underlie both resistance and suppression in our E2 clones. We performed targeted metabolic profiling on the parental parasite strain as well as E2 clones R1-R3 and S1-S2 (Fig. 7A, Table S1). We find that levels of the MEP pathway intermediate DOXP are significantly increased in FSM^R^ (*had2*^*R157X*^, *PFK9*^*WT*^) strains (Fig. 7A, p≤0.05, one-way ANOVA and Sidak’s post-test). FSM is competitive with DOXP for inhibition of its target enzyme DXR. Therefore, our data indicate that the FSM^R^ strains achieve resistance via increased levels of DOXP, which outcompetes FSM. We also observe a significant increase in the downstream MEP metabolite, MEcPP, in our FSM^R^ strains (p≤0.05).

**Figure 7.**
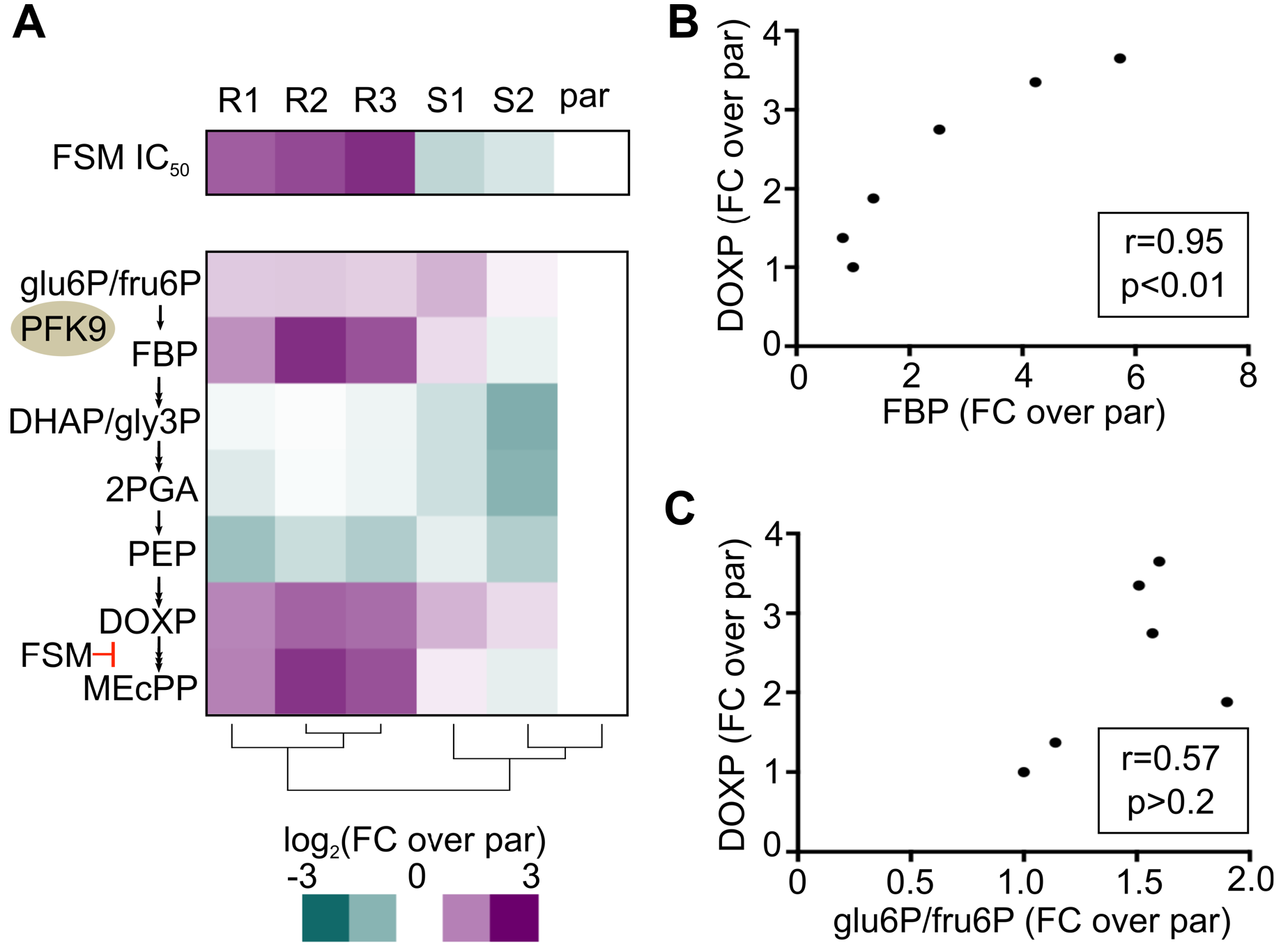
*HAD2* and *PFK9* alleles alter FSM resistance and metabolite levels in *P. falciparum*. **(A)** Metabolic profiling and clustering of parental and E2 clone strains demonstrates a metabolic signature of resistance, which includes increased levels of MEP pathway intermediates DOXP and MEcPP and the glycolytic metabolite and PFK product FBP. Glu6P/fru6P and DHAP/gly3P are isomer pairs that cannot be confidently distinguished. Clustering performed using the heatmap function in R. Data are also summarized in S1 Table. FSM IC_50_s are shown for reference. **(B)** DOXP levels are highly correlated to levels of the upstream glycolytic metabolic FBP (Pearson’s r=0.95). **(C)** In contrast, DOXP levels are not correlated to the glycolytic metabolites glu6P/fru6P (Pearson’s r=0.57).

To understand the role of PFK9 in conferring and suppressing FSM resistance, we determined the steady-state levels of intermediates from glycolysis, metabolites of which feed into the MEP pathway (Fig. 7A, Table S1). Hierarchical clustering indicates that resistant clones are characterized by a metabolic signature of increased levels of FBP, DOXP, and MEcPP (Fig. 7A). We observe that the abundance of DOXP and MEcPP are tightly correlated with cellular levels of the PFK9 product, FBP (Fig. 7B, p<0.01), but not with the other upstream glycolytic metabolites, such as glu6P/fru6P (Fig. 7C, p>0.2).

Of note, the *pfk9*^*T1206l*^ suppressor allele in strains S1 and S2 restores nearly parental levels of FBP and downstream MEP pathway intermediates (Fig. 7A), consistent with our finding that PFK activity is reduced in lysate from these strains (Fig. 6B).

## DISCUSSION

Cells must control levels of critical metabolites in order to efficiently utilize carbon sources for energy and biosynthesis of essential molecules. Cells may regulate their metabolism via transcriptional, post-transcriptional, post-translational, allosteric, or enzymatic mechanisms that are necessary for growth (57–60). In the glucose-rich red blood cell niche, *Plasmodium* spp. malaria parasites display a unique dependence on glycolysis for energy and biosynthesis.

Using resistance to a metabolic inhibitor, we identify a phosphatase member of the HAD superfamily, HAD2, as a novel regulator of metabolism in *P. falciparum*. Importantly, HAD2 controls substrate availability to the parasite-specific MEP pathway for isoprenoid synthesis, which is a promising drug target for much needed new antimalarials. We find that tolerance to inhibitors such as FSM is a robust and sensitive readout of metabolic perturbation. HAD2 is necessary for metabolic homeostasis in malaria parasites. Cells lacking HAD2 exhibit marked dysregulation of central carbon metabolism, including altered steady-state levels of glycolytic intermediates and isoprenoid precursors. We find that mutations in phosphofructokinase (*PFK9*) restore wild-type growth rates and FSM sensitivity to our *had2* mutant strains. Our study thus genetically connects the function of HAD2, a HAD superfamily member, to control of essential central carbon metabolism. In addition, our work reveals a previously undescribed strategy by which malaria parasites may respond to cellular metabolic dysregulation through mutation in the gene encoding the rate-limiting glycolytic enzyme PFK9.

HAD2 is a member of the HAD superfamily and a homolog of the previously described metabolic regulator HAD1. With our previous studies on HAD1 (20, 21), we define the cellular role of these proteins in *P. falciparum* and contribute to the greater understanding of the HADs, an evolutionarily conserved and widespread protein family. Both enzymes belong to the IIB (IPR006379) and Cof-like hydrolase (IPR000150) subfamilies (22). HAD enzymes display diverse substrate preferences (23, 27, 43, 44, 61–64), and their biological functions are largely unknown. Like other HAD homologs, including PfHAD1 (20), HAD2 appears to be a non-specific, cytoplasmic phosphatase with preference for small, monophosphorylated substrates. While the HAD superfamily is thought to be highly evolvable pool of enzymes with broad substrate specificity (37, 42), our work strongly suggests that, like HAD1 and HAD2, other members of this superfamily are likely to perform specific and biologically important cellular functions. As *P. falciparum* has a smaller repertoire of HADs than bacteria, and *Plasmodium* HADs influence easily quantified phenotypes (drug tolerance, growth, metabolite levels), the malaria parasite may be an attractive system for study of the molecular mechanisms by which HAD proteins control metabolic homeostasis and growth.

Metabolic profiling reveals that loss of HAD2 function leads to metabolic dysregulation, which is centered around the canonical rate-limiting step of glycolysis, catalyzed by PFK9. While the cellular abundance of the PFK9 product FBP is increased in *had2* parasites, HAD2 does not directly utilize FBP as an enzymatic substrate, suggesting an indirect mechanism of HAD2-mediated metabolic regulation. However, the distinct metabolic signature of *had2* parasites, characterized by increased levels of the MEP pathway metabolites DOXP and MEcPP and the key glycolytic metabolite FBP, suggests that MEP pathway metabolism is precisely linked to FBP production. In other microbial systems, FBP levels reflect metabolic flux and are cued to environmental perturbations (65). FBP-centered metabolic regulation is also important to the related apicomplexan *Toxoplasma gondii*, which constitutively expresses the fructose 1,6- bisphosphatase (FBPase) to fine-tune glucose metabolism (57). While *P. falciparum* does not appear to possess an FBPase, necessary for gluconeogenesis, the parasite may possess alternative FBP-sensing mechanisms to tune metabolism, perhaps via regulators such as HAD1 and HAD2.

The metabolic dysregulation we observe in the *had2* mutant strains appears to come with a fitness disadvantage. Under FSM-selective pressure, the benefits of dysregulated metabolism outweigh the costs. However, in the absence of FSM, *had2* parasites achieve metabolic relief through secondary mutation in *PFK9*. The improved growth of *had2 pfk9* double mutant parasites, compared to parasites with a *had2* single mutation, argues that both the growth and metabolic phenotypes are linked. However, our complementation studies cannot strictly discern whether restoring HAD2 directly improves the growth rate of the *had2* strain, as transfection and complementation of HAD2 is inherently an additional selection for increased fitness. Of note, two recent essentiality screens in *Plasmodium* spp. have found that HAD2 is dispensable for growth, and loss of HAD2 was not found to be associated with any significant fitness defect (66, 67). However, it is unknown whether the mutant strains generated in these screens have also acquired additional suppressor mutations, such as polymorphisms in *PFK9*, that have facilitated their growth.

Likewise, *PFK9* provides an additional case-study of the context dependence of gene essentiality in *Plasmodium* spp. As the canonical rate-limiting step in glycolysis, *PFK9* is strongly predicted to be essential for asexual growth of malaria parasites (66, 67). In the context of a *had2* mutation, our strains readily develop mutations in *PFK9* that reduce function but are nonetheless associated with increased fitness. Indeed, it is surprising that parasites that are entirely dependent on glycolysis for ATP production tolerate such a significant loss of activity in this enzyme. Because we identify mutations across the length of *PFK9* in our suppressed strains, our studies do not appear to point to a specific disrupted function, such as alterations in an allosteric binding pocket or a dimer interface. The observed mutability of *PFK9* points to a remarkable and unexpected metabolic plasticity in the parasite. That is, even though the parasite inhabits a highly controlled intraerythrocytic niche, a wide range of metabolic states of *P. falciparum* growth are still permissive for parasite growth. This previously undescribed metabolic plasticity centered on PFK9 should be considered in future efforts to target essential metabolism in *Plasmodium*.

Combined with the above study, our work highlights the central role of the glycolytic enzyme PFK9. A recent kinetic model of parasite glycolysis confirms that PFK has a high flux-control coefficient, is sensitive to competitive inhibition, and can effectively reduce glycolytic flux (68, 69). Like HAD2, PFK9 is plant-like and evolutionarily divergent from its mammalian homologs (51). These differences may be exploited for PFK inhibitor design and may indicate that PFK9 can be specifically targeted for antimalarial development. However, our work cautions that the parasite has a surprising capacity to adapt to perturbations in central carbon metabolism, which may present challenges when targeting these pathways.

Finally, our approach demonstrates the power of forward genetics to uncover novel biology in a clinically relevant, non-model organism. We employ a previously described screen (20) to uncover a novel resistance locus and employ a second selection for parasite fitness to identify changes that suppress our resistance phenotype. Of the 19 strains in our original FSM resistance screen (20), we identify only one *had2* mutant, likely due to the reduced fitness associated with resistance in this strain. While fitness costs associated with antimalarial resistance are well known (70–73), this study represents, to our knowledge, the first to harness this evolutionary trade-off to identify suppressor mutations in a non-target locus. Additional methods to identify low fitness resistant mutants have recently been recently described (73), and fitness assessment of resistance mutations may allow for suppressor screening for other antimalarials or other target pathways to reveal new aspects of biology and drug resistance in malaria parasites.

## MATERIALS AND METHODS

### Parasite strains and culture

Unless otherwise indicated, parasites were maintained at 37 °C in 5% O_2_/5% CO_2_/90% N_2_ in a 2% suspension of human erythrocytes in RPMI medium (Sigma Aldrich) modified with 27 mM NaHCO_3_, 11 mM glucose, 5 mM HEPES, 0.01 mM thymidine, 1 mM sodium pyruvate, 0.37 mM hypoxanthine, 10 μg/mL gentamycin, and 5 g/L Albumax (Thermo Fisher Scientific).

FSM^R^ strain E2 was generated by selecting a clone of genome reference strain 3D7 (MRA-102, MR4, ATCC, Manassas, VA) under continuous treatment with 3 μM FSM, as previously described (20). Clones of strain E2 were isolated by limiting dilution.

### Quantification of FSM resistance

Opaque 96-well plates were seeded with asynchronous cultures at 0.5-1.0% parasitemia (percent of infected red blood cells). After 3 days, media was removed and parasitemia was measured via Picogreen fluorescence on a POLARStar Omega spectrophotometer (BMG Labtech), as previously described (74). Half maximal inhibitory concentration (IC_50_) values were calculated using Graphpad Prism. Unless otherwise indicated, all IC_50_ data are representative of means from at least ≥3 independent experiments performed with technical replicates.

### HAD2 structural alignment

Structures were aligned using the TM-align algorithm in Lasergene Protean 3D software (root mean square deviation (RMSD) of 1.9 Å).

### Whole genome sequencing and variant analysis

Library preparation, sequencing, read alignment, and variant analyses were performed by the Washington University Genome Technology Access Center. One microgram of parasite genomic DNA was sheared, end repaired, and adapter ligated. Libraries were sequenced on an Illumina HiSeq 2500 in Rapid Run mode to generate 101 bp paired end reads. Reads were aligned to the *P. falciparum* 3D7 reference genome (PlasmoDB v7.2) using Novoalign (V2.08.02). Duplicate reads were removed. SNPs were called using samtools (quality score ≥20, read depth ≥5) and annotated using snpEff. Background variants were removed using previously sequenced genomes from parental and control strains (20). Mixed variant calls and variants in highly variable surface antigen loci (75, 76) were removed.

### Sanger sequencing

The *HAD2* (PlasmoDB PF3D7_1226300) A469T (R157X) SNP was amplified and sequenced using the HAD2_R157X_F and HAD2_R157X_R primers. The *PFK9* locus was amplified using the PFK9_F and PFK9_R primers. *PFK9* amplicons were sequenced using the PFK9_seq (1-8) primers. Primer sequences can be found in Table S2.

### Generation of recombinant HAD2

The predicted coding sequence of *HAD2* was amplified using the HAD2_LIC_F and HAD2_LIC_R primers (Table S2). A catalytic mutant (D26A) was also generated to use as a negative control in activity assays. The *had2*^*D26A*^ allele was created using the HAD2_D26A_F and HAD2_D26A_R site-directed mutagenesis primers (Table S2).

Ligation-independent cloning was used to clone *HAD2* and *had2*^*D26A*^ into vector BG1861 (77), which introduces an N-terminal 6xHis fusion to the expressed protein. BG1861:6xHis-HAD2 was transformed into One Shot BL21(DE3)pLysS *Escherichia coli* cells (Thermo Fisher Scientific). Protein expression was induced for 3 hours with 1 mM isopropyl-β-D-thiogalactoside at mid-log phase (OD_600_ 0.4-0.5). Cells were collected by centrifugation and stored at −20°C.

Cells were lysed in buffer containing 1 mg/mL lysozyme, 20 mM imidazole, 1 mM dithiothreitol, 1 mM MgCl_2_ 10 mM Tris HCl (pH 7.5), 30 U benzonase (EMD Millipore), and EDTA-free protease inhibitor tablets (Roche). 6xHis-HAD2 was bound to nickel agarose beads (Gold Biotechnology), washed with 20 mM imidazole, 20 mM Tris HCl (pH 7.5), and 150 mM NaCl and eluted in 300 mM imidazole, 20 mM Tris HCl (pH 7.5), and 150 mM NaCl. This eluate was further purified by size-exclusion gel chromatography using a HiLoad 16/600 Superdex 200 pg column (GE Healthcare) equilibrated in 25 mM Tris HCl (pH 7.5), 250 mM NaCl, and 1 mM MgCl_2_. The elution fractions containing HAD2 were pooled and concentrated and glycerol was added to 10% (w/v). Protein solutions were immediately flash frozen and stored at −80°C.

### HAD2 activity assays

The rate of *para*-nitrophenyl phosphate (*p*NPP; Sigma-Aldrich S0942) hydrolysis by HAD2 was determined by continuous measurement of absorbance at 405 nm. Assays were performed at 37°C in 50 μl with 50 mM Tris-HCl, pH 7.5, 5 mM MgCl_2_, 10 mM *p*NPP, and 1.2 μM enzyme.

Hydrolysis of other phosphorylated substrates by HAD2 was measured using the EnzChek phosphate assay kit (Life Technologies). Reaction buffer was modified to contain 50 mM Tris-HCl, 1 mM MgCl_2_, pH 7.5, 0.1 mM sodium azide, 0.2 mM 2-amino-6-mercapto-7-methylpurine riboside (MESG), and 1 U/mL purine nucleoside phosphorylase (PNP). Reactions contained 5 mM substrate and 730 nM enzyme. Activity was normalized to that obtained from catalytically inactive 6xHis-HAD2^D26A^. All data are means from ≥3 independent experiments performed with technical replicates.

### *P. falciparum* growth assays

Asynchronous cultures were seeded at 1% parasitemia. Media (no drug) was exchanged daily. Samples were taken at indicated timepoints and fixed in PBS + 4% paraformaldehyde, 0.05% glutaraldehyde. Cells were stained with 0.01 mg/ml acridine orange and parasitemia was determined on a BD Biosciences LSRII flow cytometer (Thermo Fisher Scientific). All data are means from ≥3 independent experiments.

### pTEOE110:HAD2 plasmid construction

The pTEOE110 construct contains the *heat shock protein 110* (PF3D7_0708800) 5’ UTR and a C-terminal GFP tag (20). Human dihydrofolate reductase (hDHFR) is present as a selectable marker. Inverted terminal repeats are included for genome integration by a co-transfected piggyBac transposase (pHTH, MRA912, MR4, ATCC, Manassas, VA).

*HAD2* was amplified with the HAD2_XhoI_F and HAD2_AvrII_R primers (Table S2) and cloned into AvrII and XhoI sites in the pTEOE110 plasmid.

### Parasite transfections

Transfections were performed as previously described (20). Briefly, 50-100 μg of plasmid DNA was precipitated and resuspended in Cytomix (25 mM HEPES pH 7.6,
120 mM KCl, 0.15 mM CaCl_2_, 2 mM EGTA, 5 mM MgCl_2_, 10 mM K_2_HPO_4_).

A ring-stage *P. falciparum* culture was washed with Cytomix and resuspended in the DNA/Cytomix solution. Cells were electroporated using a BioRad Gene Pulser II electroporator at 950 μF and 0.31 kV. Electroporated cells were washed with media and returned to normal culture conditions. Parasites expressing the construct were selected by continuous treatment with 5 nM WR92210 (Jacobus Pharmaceuticals). Transfectants were cloned by limiting dilution and presence of the HAD2-GFP construct was verified by PCR using gene- and GFP-specific primers (HAD2_R157X_F and GFP_R, Table S2). Maintenance of the endogenous *HAD2* and *PFK9* genotypes were verified by Sanger sequencing.

### Antiserum generation

Polyclonal anti-HAD2 antiserum was raised in rabbits against 6xHis-HAD2, with Titermax as an adjuvant (Cocalico Biologicals). Antiserum specificity was confirmed by immunoblot of lysate lacking HAD2. Polyclonal anti-HAD1 antiserum has been previously described (MRA-1256, MR4, ATCC) (20).

### Immunoblotting

Lysates were separated on a polyacrylamide gel and transferred to a polyvinylidene difluoride membrane. Membranes were blocked in 5% non-fat dry milk, 0.1% Tween-20 in PBS. Rabbit polyclonal antisera were used at the following dilutions: 1:2,000-5,000 anti-HAD2 and 1:20,000 anti-HAD1 (20). For all blots, 1:20,000 HRP-conjugated goat anti-rabbit IgG antibody was used as a secondary antibody (ThermoFisher 65-6120). Blots were stripped with 200 mM glycine, 0.1% SDS, 1% Tween-20, pH 2.2 and reprobed with 1:5,000 rabbit anti-heat shock protein 70 (Hsp70) (AS08 371, Agrisera Antibodies) as a loading control. All blots shown are representative of ≥3 independent experiments. Minimal adjustments were applied equally to all blot images.

### PfPFK model construction

PfPFK subunits were searched against the HHpred server for protein remote homology detection and 3D structure prediction using statistics as previously described (78–81). The *Borellia burgdorferii* PFK structure (PDB 1KZH) (82) returned the highest similarity for both PfPFK domains and was used to predict 3D structure for each domain using the program MODELLER. PFK product orientation in the active site of the model was predicted via the alignment tool, using PyMOL software against the *E. coli* PFK crystal structure (PDB 1PFK) (83). The α domain model encompasses amino acids 779-1347 and the β domain model encompasses amino acids 110-638.

### Assay of native PFK9 activity

Sorbitol-synchronized trophozites were isolated using 0.1% saponin. Cells were washed in buffer containing 100 mM Tris-HCl (pH 7.5), 1 mM MgCl_2_, 1 mM DTT, 10% glycerol, and EDTA-free protease inhibitor tablets (Roche) and lysed by sonication at 4°C (Fisher Scientific Model 550 Sonic Dismembrator, amplitude 3.5), followed by centrifugation at 4°C (10,000 × *g*, 10 min). An “RBC carryover” control was comprised of the trace cellular material remaining after saponin lysis, centrifugation, and wash of uninfected erythrocytes.

Lysate PFK9 activity was monitored by linking it to the oxidation of NADH, as previously described (51, 56). Reactions contained 100 mM Tris-HCl (pH 7.5), 1 mM MgCl_2_, 1 mM DTT, 0.25 mM NADH, 1 mM ATP, 3 mM fructose 6-phosphate, and excess of linking enzymes aldolase (7.5 U), triose-phosphate isomerase (3.8 U), and glycerol 3-phosphate dehydrogenase (3.8 U). After adding fresh cell lysate (10-15 μg total protein), absorbance at 340 nm was measured at 37°C for 40 min. Activity was determined by linear regression using Graphpad Prism software. Unless otherwise indicated, data are means from ≥3 independent experiments.

### Metabolite profiling

Approximately ~1 × 10^9^ sorbitol-synchronized early trophozites were isolated using 0.1% saponin, washed with ice-cold PBS + 2 g/L glucose, and frozen at −80 °C. Samples were extracted in 600 μL of ice-cold extraction solvent [chloroform, methanol, and acetonitrile (2:1:1, v/v/v)] using two liquid-nitrogen cooled 3.2 mm stainless steel beads and homogenization in a Tissue-Lyser II instrument (Qiagen) at 20 Hz for 5 minutes in a cold sample rack. Ice-cold water was added and samples were homogenized for 5 min at 20 Hz. Samples were centrifuged at 14,000 rcf at 4°C for 5 min. The polar phase was lyophilized and re-dissolved in 100 μL water and analyzed by LC-MS/MS. LC-MS/MS was performed on a 4000QTRAP system (AB Sciex) in multiple-reaction monitoring mode using negative ioniziation and 10 mM tributylammonium acetate (pH 5.1-5.5) as the ion pair reagent. The specific parameters used for analysis of MEP pathway metabolites have been previously described (19). Liquid chromatography separation was performed using ion pair reverse-phase chromatography (84) with the following modifications: (1) RP-hydro 100 mm × 2.0 mm, 2.5 μm high performance liquid chromatography column (Phenomenex), (2) flow rate of 0.14 mL/min, (3) solvent A of 10 mM tributylammonium acetate in 5% methanol, (4) binary LC gradient (20% solvent B (100% methanol) from 0 to 2.5 min, 30% B for 12.5 min, 80% B for 5 min, and column equilibration at for 5 minutes), and (5) 20 μL autosampler injection volume.

### Accession numbers

All genome data have been deposited in the NCBI BioProject database (PRJNA222697) and Sequence Read Archive (SRP038937).

## ACKNOWLEDGEMENTS

We thank the Genome Technology Access Center in the Department of Genetics at Washington University School of Medicine for help with genomic analysis. The Center is partially supported by NCI Cancer Center Support Grant #P30 CA91842 to the Siteman Cancer Center and by ICTS/CTSA Grant# UL1TR002345 from the National Center for Research Resources (NCRR), a component of the National Institutes of Health (NIH), and NIH Roadmap for Medical Research. This publication is solely the responsibility of the authors and does not necessarily represent the official view of NCRR or NIH. We thank Sophie Alvarez and the Proteomics and Mass Spectrometry Facility at the Donald Danforth Plant Science Center for performing the LC-MS/MS sample prep and analysis. This work was supported by the National Science Foundation under grant number DBI-0521250 for acquisition of the QTRAP LC-MS/MS. We thank Daniel Goldberg (Washington University) for supplying the pTEOE110 plasmid. We thank the Malaria Research Reference and Reagent Resource (MR4) for providing reagents contributed by D.J. Carucci (MRA-102) and John Adams (MRA-912). We thank Katherine Andrews for sharing unpublished data relevant to this study.

## SUPPLEMENTAL MATERIAL LEGENDS

**Supplemental Figure 1. FSM^R^ strain E2 does not have increased expression of *DXS*, *DXR*, or *HAD1*.**

**Supplemental Figure 2. HAD2 is cytosolic.**

**Supplemental Figure 3. Assay of PFK activity from *P. falciparum* lysate is linear and specific.**

**Supplemental Table 1. Relative metabolite levels in strains described in this study.**

**Supplementary Table 2. Primers used in this study.**

